# The immunogenic 33-mer gliadin peptide attenuates flagellin-induced TLR5/NF-κB signaling

**DOI:** 10.64898/2026.10.10.758210

**Authors:** Karen M. Lammers, Diego S. Vazquez, Veronica Segura, Jürgen Hollmann, Wolfgang Hübner, Servaas A. Morré, Thomas Huser, Carolina Sousa, Veronica I. Dodero

**Affiliations:** Department of Chemistry, Bielefeld University, Universitätsstr. 25, 33615 Bielefeld, Germany; Laboratorio de Expresión y Plegado de Proteínas (LEPP), Departamento de Ciencia y Tecnología, Universidad Nacional de Quilmes, Roque Sáenz Peña 352 (B1876BXD), Bernal, Buenos Aires, Argentina; Grupo de Biología Estructural y Biotecnología (GBEyB-IMBICE), Departamento de Ciencia y Tecnología, Universidad Nacional de Quilmes, Roque Sáenz Peña 352 (B1876BXD), Bernal, Buenos Aires, Argentina; Consejo Nacional de Investigaciones Científicas y Técnicas (CONICET), Av. Rivadavia 1917 (C1033AAJ), Ciudad Autónoma de Buenos Aires, Argentina; Department of Microbiology and Parasitology. Faculty of Pharmacy. University of Seville. Seville, Spain; Faculty of Physics, Biomolecular Photonics, Bielefeld University, Universitätsstr. 25, 33615 Bielefeld, Germany; Institute for Public Health Genomics (IPHG), Department of Genetics and Cell Biology, Research School GROW (School for Oncology & Developmental Biology), Faculty of Health, Medicine & Life Sciences, University of Maastricht, Universiteitssingel 40, 6229 ER Maastricht, The Netherlands

## Abstract

Wheat gluten is the main trigger of gluten-related disorders such as celiac disease, in which the proteolysis-resistant, 33-mer peptide is a key immunodominant factor. It has been reported that the 33-mer forms oligomers and thin-plate superstructures that accumulate in the Caco-2 intestinal epithelial cell line, although their biological effects remain only partially understood. Here, we screened the effects of these nanostructures on Toll-like receptors (TLRs) in Caco-2 cells to investigate their effect on the early innate immune response. Our findings showed a specific upregulation of TLR5 mRNA in response to the 33-mer gliadin system, although no inflammatory response was detected based on CXCL1 and CXCL8 levels. Moreover, immunofluorescence analysis revealed partial colocalization of 33-mer with TLR5. Finally, we found that 33-mer attenuates flagellin-induced TLR5/NF-κB signaling in a concentration-dependent manner in reporter cells, similar to a blocking anti-hTLR5-IgA2 antibody. Molecular docking suggested that the 33-mer peptide could occupy the concave region of TLR5 and 54% of the TLR5– TLR5 dimerization interface. Altogether, our findings reveal a previously unrecognized capacity of the 33-mer to modulate flagellin/TLR5/NF-κB signaling pathway in model systems. We hypothesize that 33-mer supramolecular nanostructures may act as local reservoirs, increasing local 33-mer concentration and potentially facilitating its interaction with TLR5, consistent with a plausible binding mode suggested by molecular docking studies. Because TLR5 contributes not only to inflammation but also to intestinal homeostasis, these findings warrant further investigation into how 33-mer supramolecular organization modulates TLR5 signaling as part of the early innate immune response to gluten peptides.

## Introduction

Gluten proteins from wheat, barley, rye, and some oats are only partially digested by human digestive proteases (1,2). As a result, large proteolysis-resistant gluten peptides of 20-40 amino acids reach the intestinal epithelium and enter the gut mucosa (3), triggering gluten-related disorders (GRD) in predisposed individuals (4–6). Among GRD, Celiac disease (CeD) is the autoimmune presentation, a common gastrointestinal disorder with an estimated global prevalence of up to 1%, whose adaptive immune response has been studied but whose early steps of immune activation remain unsolved (6–9).

Among proteolysis-resistant gliadin fragments, the 33 amino acid α-gliadin peptide (33-mer) is particularly relevant because of its central role in the adaptive immune response associated with CeD (10). Beyond its well-established immunogenicity, the known structural and supramolecular properties of the 33-mer raise intriguing questions about potential biological functions that remain largely unexplored (11). Recently, it was reported that the 33-mer forms different-sized oligomers that accumulate in the gut epithelial cell model Caco-2 without impairing the barrier function (12) and without exhibiting pro-inflammatory activity, unlike pepsin-digested gliadin nanostructures (13). From the molecular point of view, the 33-mer gliadin peptide is intrinsically disordered - a peptide sequence lacking a globular folded structure and behaving as a highly dynamic and flexible conformation - whose major structural motif is the polyproline II (PPII) helix secondary structure and exists as a monomer, coexisting with small oligomers and superstructures (**Figure 1**) (12,14–19). At concentrations where the 33-mer forms thin-plate nanostructures, activation of Toll-like Receptors TLR4 and TLR2 was observed in macrophages; however, other TLRs have not yet been investigated (16). In general and particularly in CeD, TLR signaling is investigated in professional immune cells, such as macrophages and dendritic cells (20, 21). However, its role in polarized intestinal epithelial monolayers remains less explored. Given the recurrent in vivo exposure of intestinal epithelial cells to proteolysis-resistant gluten peptides, whether the accumulation and supramolecular organization of the 33-mer contribute to the modulation of epithelial TLR signaling remains to be elucidated.

**Figure 1:**
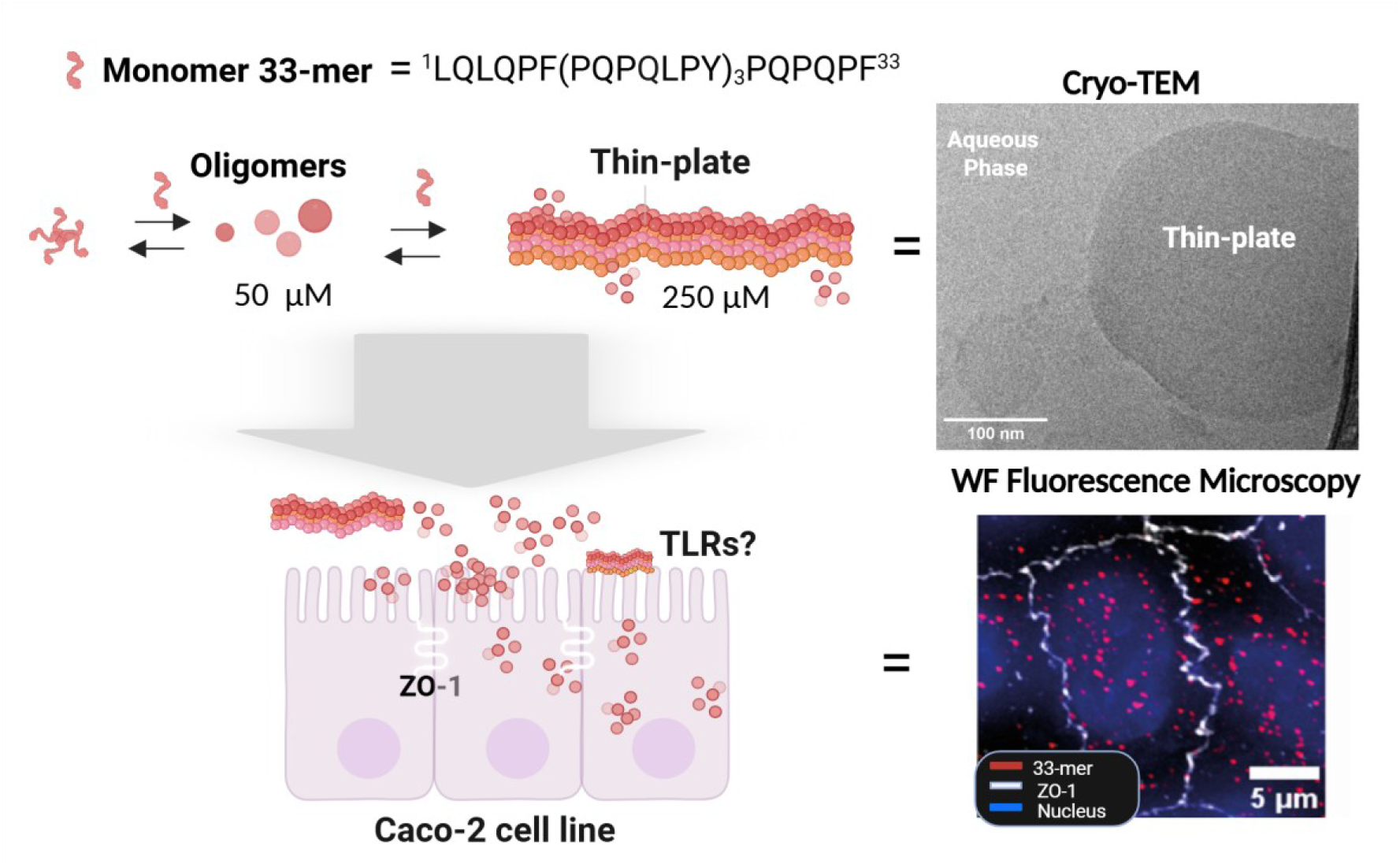
Schematic representation of the 33-mer self-assembly process towards the formation of thin-plate superstructures detected previously by Cryo-TEM (17) and 33-mer accumulation (red) in differentiated monolayers of the Caco-2 cell line maintaining the barrier function (intact Zonula occludens 1 (ZO-1, in white) (12).

Based on that, the aim of this study was to investigate the effect of the 33-mer gliadin peptide supramolecular system on TLR-related responses in Caco-2 monolayers. We first examined TLR mRNA expression and inflammatory responses in differentiated Caco-2 monolayers exposed to 33-mer at concentrations corresponding to different supramolecular assemblies, including oligomers and thin-plate nanostructures (**Figure 1**). Guided by this initial screening, we further investigated whether the 33-mer could modulate TLR5/NF-κB signaling using a TLR5 reporter-cell system. Finally, molecular docking was employed to explore a possible structural basis for the observed functional effect of the 33-mer system on the flagellin/TLR5/NF-κB signaling pathway. Together, these approaches aim to shed light on the relationship between the supramolecular organization of gliadin peptides and TLR-mediated innate immune signaling.

## Methods

### Reagent and sample preparation

The 33-mer gliadin peptides and their oligomers were obtained and characterized as previously described (12,14–16). Briefly, The 33-mer (or 33-mer DGP) was dissolved in PBS buffer (10 mM phosphate buffer, 2.7 mM potassium chloride, and 137 mM sodium chloride, pH 7.4 at 25 °C, Aldrich P4417) at a final concentration of 766 µM as a stock solution, followed by stabilizing the 33-mer colloidal system by incubating it three days at 4 °C. P-gliadin was prepared by pepsin digestion of gliadin (Aldrich), using a previously described protocol (13). Ultrapurified flagellin from *S. typhimurium* and neutralizing anti-hTLR5-IgA2 were purchased from Invivogen (France). Sodium butyrate was purchased from Sigma-Aldrich (USA). The monoclonal antibody (mAb) mouse anti-33-mer (G12) was purchased from Biomedal (Spain), and the rabbit anti-TLR5 antibody was purchased from Invitrogen (Germany).

### Cell culture

Enterocytes (passage 18-40) from the Caco-2 cell line (ATCC, UK) were cultured in 75 cm^2^ flasks (Falcon, Corning Incorporated, USA) using a culture medium consisting of Minimal Essential Medium supplemented with 10 % heat-inactivated fetal bovine serum, 2 mM L-glutamine, 1 % non-essential amino acids (Gibco, USA) and 1 % penicillin/streptomycin (Sigma-Aldrich, USA) at 37 °C in a 5 % CO2 incubator. Cells were passaged at 80% confluency. For stimulation experiments, Caco-2 enterocytes were seeded in 12-well plates at 3.8 × 10^5^ cells/well and in 6-well plates at 9.5 × 10^5^ cells/well (Costar, Corning Incorporated). Cells were grown for 12 days after seeding to reach confluence. The alkaline phosphatase (ALP) assay showed that after 12 days of cultivation, the ALP signal reached a plateau, suggesting monolayer differentiation. For transport and microscopy experiments, the Caco-2 cells were grown on 12-well culture inserts (Sarstedt, pore size 0.4 µm, growth area 1.13 cm^2^) for 21 days. The medium was changed three times a week. For the assessment of TLR expression and the production of chemokines, monolayers of Caco-2 cells grown in 12-well plates were incubated with PBS in medium alone (PBS control) or with 33-mer suspensions at concentrations of 50 μM or 250 μM for 20 hours. In some wells, 100 ng/mL TLR5 agonist, the ultrapure flagellin from *S. Typhimurium* (Invivogen), was added to the monolayers as a positive control. As a protein control, pepsin-gliadin digest (P-gliadin) was employed. Cell viability tests are presented in the supplementary information (**SI, Table S1**). At the end of incubation, we collected supernatants and stored them at -80 °C until assay. Cells were lysed, and RNA was isolated and reverse-transcribed into cDNA for real-time RT-PCR analysis of gene expression. Supernatants were used to quantify CXCL1 and CXCL8 by DuoSet™ Enzyme-linked Immunosorbent Assay (ELISA) Development System according to the manufacturer’s protocol (R&D Systems). The detection limit for both chemokine assays was 31.3 pg/mL. For transport and microscopy experiments, transepithelial electrical resistance (TEER) was measured before and after stimulation (**SI, Fig.S2**). Basolateral concentrations of 33-mer were measured by ELISA assays with mAb G12 (Biomedal).

### Real-time PCR

The RNA was extracted from Caco-2 enterocytes with the RNeasy kit (Qiagen, USA). Complementary DNA (cDNA) was synthesized from 1 µg RNA per sample using the RevertAid First Strand cDNA Synthesis kit (ThermoFisher Scientific, Germany). Real-time reverse transcription-polymerase chain reaction (RT-PCR) was used. Messenger RNA (mRNA) expression for an array of primers was quantified by RT-PCR using SYBR Green fluorescence (Applied Biosystems, Foster City, CA). Specific primers, purchased from Eurofins Genomics (Ebersberg, Germany), were:

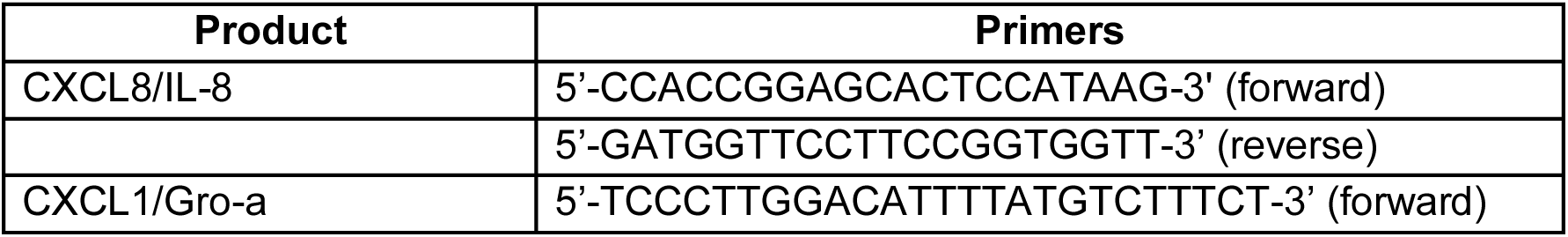
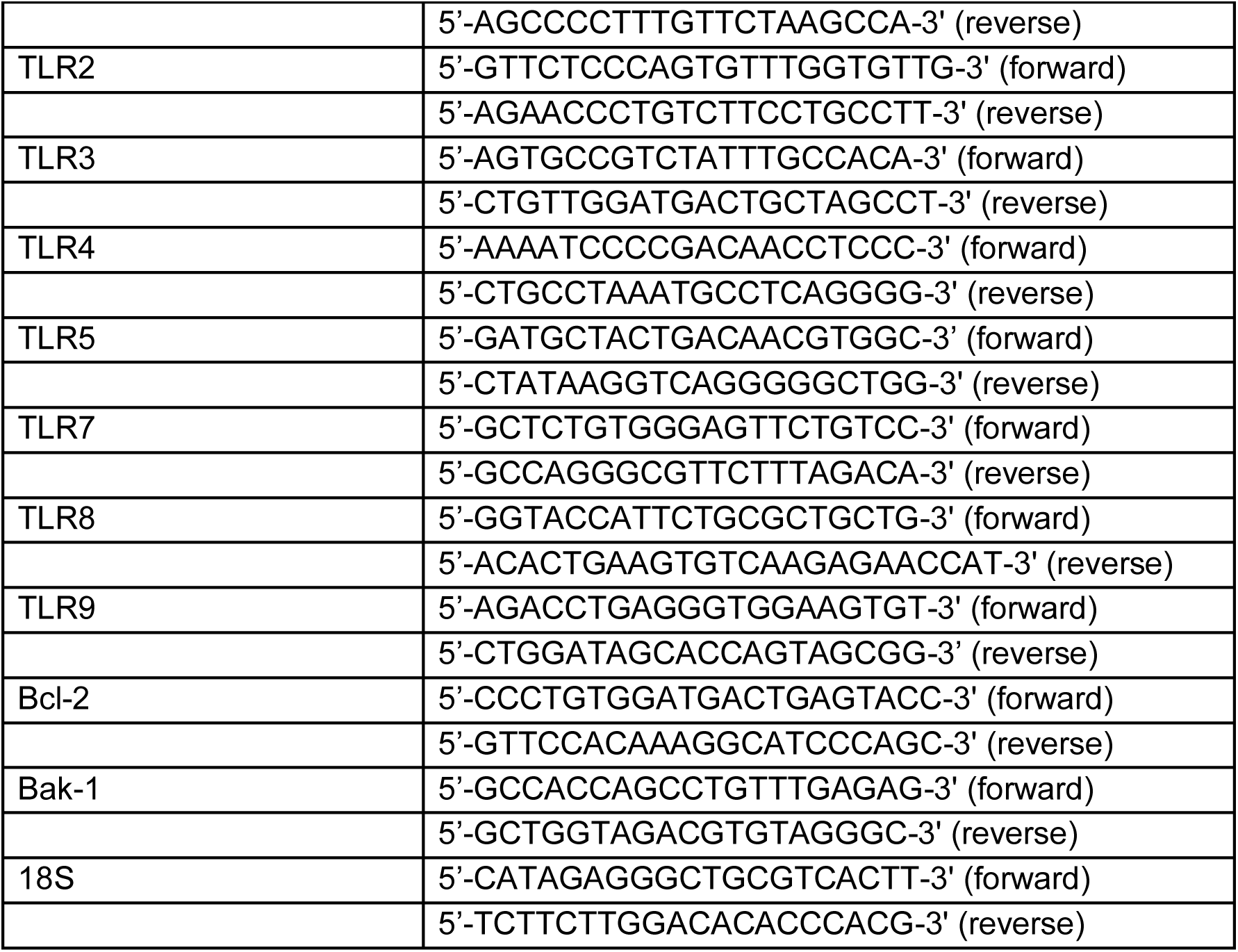

The cDNA was run on a LightCycler^®^ 480 System (Roche Molecular Diagnostics, Pleasanton, CA) using the following protocol: 1 cycle at 95 °C for 3 minutes; 45 cycles of 10 seconds at 95 °C, 15 seconds at 60 °C, and 20 seconds at 72 °C. We calculated relative gene expression using the ΔCt method, with 18S as the housekeeping gene. Relative gene expression was calculated as 2^-ΔCt^. The fold change in gene expression after stimulation was relative to the PBS control (normalized to 1). Values are presented as the fold ± standard error of the mean (SEM).

### Cell viability

To assess cell viability after treatment, we calculated the percentages of live and dead cells by Trypan blue exclusion (SI, Table S1a). In addition, the lactate dehydrogenase (LDH) release was measured as a means of cell stress and damage and compared it with a positive control (H2O-lysed cells). The results were expressed as milliunits of LDH activity per milligram of protein assayed (**Figure 2C**). In a third assay, apoptosis was determined by calculating the ratios of Bak-1 (pro-apoptotic) and Bcl-2 (anti-apoptotic) gene expression. As a moderate positive control for the induction of apoptosis in Caco-2 cells, 20 mM sodium butyrate (23) was added to parallel monolayers (SI, Table S1b).

**Figure 2:**
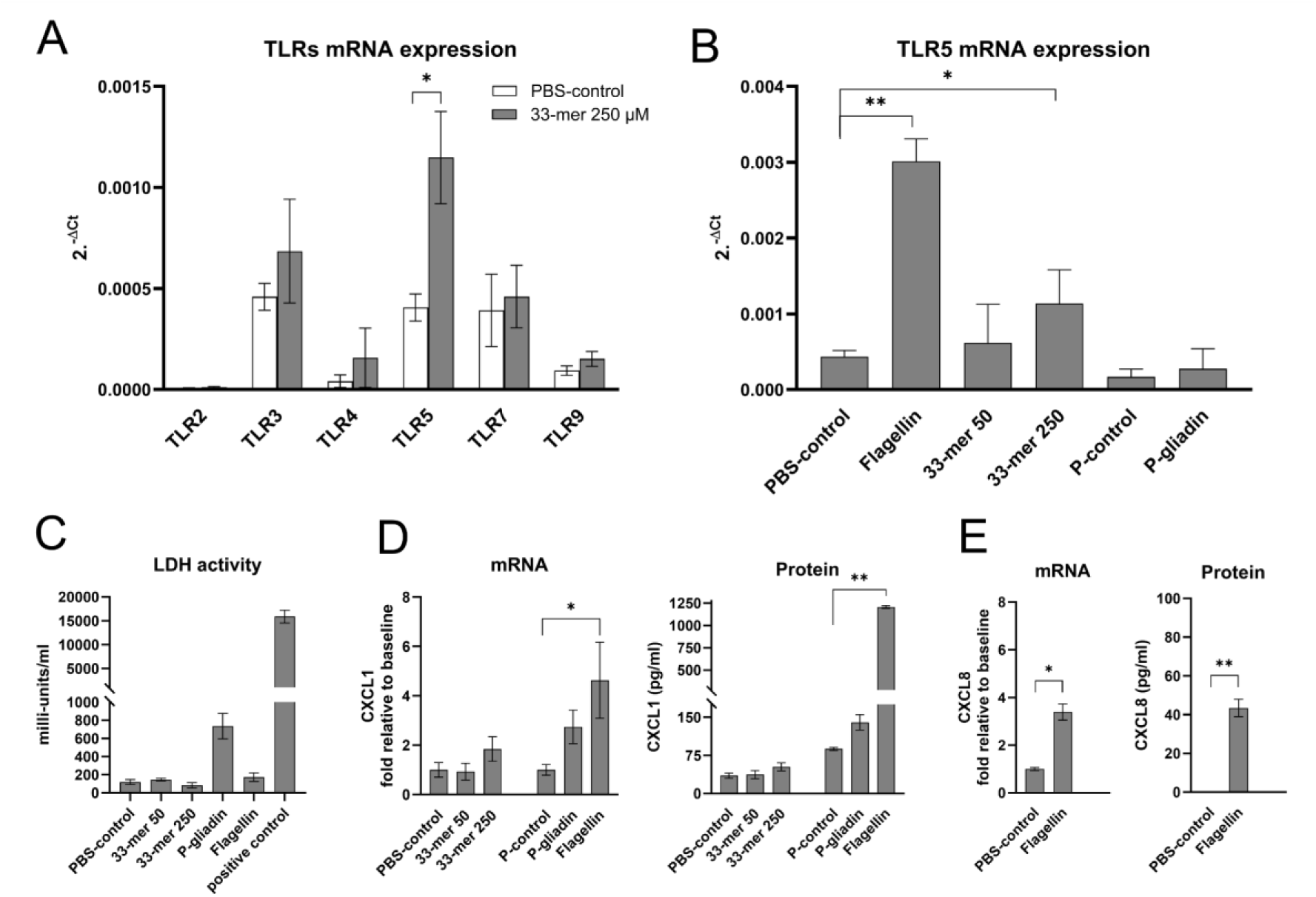
The 33-mer peptide selectively increases TLR5 gene expression in Caco-2 monolayers without damaging the cell or triggering significant CXCL1 production. (A) Screening of TLR expression in Caco-2 cells revealed low to moderate expression of TLR3, TLR4, TLR5, TLR7 and TLR9 at baseline, but only TLR5 expression was upregulated after stimulation with 33-mer at a concentration of 250 µM (*p*<0.05). (B) Flagellin, the natural ligand for TLR5, was used as a positive control. At a concentration of 100 ng/mL, flagellin induced high TLR5 gene expression in Caco-2 cells. The gliadin digest (P-gliadin) did not upregulate TLR5 gene expression. (C) To measure stimulus-induced cellular damage, supernatants from all cellular experiments were assessed for LDH activity, showing virtually no, or in the case of P-gliadin, very low, release of LDH compared to the positive control of the assay. (D) Caco-2 monolayers produced moderate levels of CXCL1 at baseline. Compared to unstimulated cells, incubation with the highest concentration of 33-mer (250 μM) increased CXCL1 production without reaching significance (P=NS). Stimulation with flagellin (100 ng/mL) increased CXCL1 gene expression (*p*<0.05) and induced high titers of CXCL1 (*p*<0.01). (E) Flagellin stimulation significantly increased CXCL8 mRNA expression (p < 0.05) and CXCL8 protein production (p < 0.01). Data are presented as fold ± SEM (n = 4) for gene expression and as mean ± SEM for protein production (n = 4). Statistical significance is indicated as p < 0.05 (*) and p < 0.01 (**).

### Transport experiments and wide-field deconvolution microscopy

Cells grown on 12-well culture inserts were differentiated and stimulated with PBS control or 250 μM 33-mer for 20 hours. At the end of incubation, the cells were fixed and prepared for fluorescence microscopy to visualize the 33-mer and TLR5. Immunofluorescence imaging, paired with deconvolution wide-field microscopy, was used to image fixed and immunostained Caco-2 cells. For this purpose, we used an inverted wide-field microscope from Deltavision (Cytiva, Marlborough, MA, USA) with a CoolSnapHQ2 CCD camera and an Olympus UplanSApo 60x water immersion objective. To visualize the 33-mer, we used the monoclonal mouse anti-G12 antibody (G12 mAb, Biomedal). This antibody is commonly used in immunoassays to quantify immunogenic gluten peptides in human fecal and urine samples (24–30). It has also been used to detect the 33-mer in biopsies from CeD patients (31,32) and Caco-2 cells (12). As the secondary antibody, goat anti-mouse Alexa Fluor 647 Ab (red) was used. TLR5 was stained with a polyclonal rabbit anti-TLR5 Ab (Invivogen) and, as a secondary antibody, a goat anti-rabbit Alexa-568 Ab (green). Colocalization of the 33-mer and TLR5 appeared in yellow. DAPI was used for staining the nuclei (blue). Microscopy images were visualized, processed, and prepared for presentation using the Open Microscopy Environment software (OMERO) (33). The methodology for calculating colocalization is described in SI (Section 5). Briefly, for representative areas where signals from the fluorescence-labeled 33-mer and TLR5 overlap, we calculated colocalization using the ImageJ plugin Coloc 2. Twelve representative areas were outlined using the freehand tool, and Coloc2 was used to analyze colocalization between the two signals across the image stack (34). Values were determined, and the mean and corresponding standard deviation were calculated. A Pearson Correlation Coefficient (PCC) tending toward 1 indicates a highly linear relationship between the pixels of the two images.

### NF-κB activation assay

HEK-Blue™-hTLR5 (Invivogen) cells are stably transfected and overexpress human TLR5 (35). HEK-Blue™-Null cells were used as a control cell line. Upon NF-κB activation, HEK-Blue™-hTLR5 cells express secreted embryonic alkaline phosphatase (SEAP), whose enzymatic activity can be quantified spectrophotometrically using HEK-Blue™ detection medium (InvivoGen, catalog no. hb-det3), according to the manufacturer’s instructions (35). To assess the specificity of TLR5-mediated signaling, a neutralizing anti-hTLR5-IgA2 antibody (0.5 µg/mL) was used to inhibit TLR5 activation induced by flagellin (10 ng/mL; InvivoGen).

### TLR5/33-mer molecular docking

The crystal structure of the TLR5/flagellin complex (PDB ID: 3V47, chain A, herein TLR5 in green) (36) was used as a template for docking. We performed blind docking using the HDock web server (37). Two systems were prepared: one TLR5 with the monomeric 33-mer peptide (red), and the other TLR5 with the monomeric deamidated 33-mer DGP peptide (blue), all previously obtained as an equilibrated structure by molecular dynamics simulation (19). The docking scores were calculated using the knowledge-based iterative scoring function ITScore PP (Protein–Protein) implemented in the HDock server. Following the empirical formula established in the HDock framework, docking scores above 0.70 indicate a high probability of binding (37). We used the highest-scoring cluster with a confidence score > 0.90 for each system for further analysis and visualization in VMD 1.9.3 (38). Contact analysis was performed using an *ad hoc* Biopython (39) script with a cutoff of 3.5 Å, and calculated solvent-accessible surface area (SASA) using the Shrake-Rupley algorithm.

### Statistical analysis

Mann-Whitney *U* tests were used to test differences between two groups. Differences among more than two groups were assessed using the Kruskal-Wallis test and Dunn’s multiple comparisons test. Values of *p*<0.05 were regarded as significant.

## Results and Discussion

### The 33-mer peptide increases the expression of TLR5

To investigate the effects of the 33-mer gliadin peptide and its supramolecular assemblies on intestinal epithelial TLR responses, we employed differentiated Caco-2 monolayers, a well-established model of the intestinal epithelial barrier (40). First, we assessed epithelial TLR gene expression and stimulus-induced changes in response to 33-mer at two selected concentrations, 50 µM and 250 µM, as plausible low- and high-dose exposures, corresponding to monomers in equilibrium with oligomers and thin-plate nanostructures, respectively (14–17). These concentrations were selected based on reported levels of habitual gluten consumption (5-20 g/day) and the cultivar-dependent abundance of potentially releasable 33-mer in wheat flour (91–603 µg/g flour) (41), while recognizing that they represent controlled *in vitro* exposures rather than direct estimates of intestinal concentrations *in vivo* (**Figure 1**). Additionally, we employed P-gliadin, the product of pepsin-mediated gliadin proteolysis, as a control, which is known to form amyloid-like fibrils and oligomers that activate mRNA expression of markers involved in recruiting immune cells (13).

Under our experimental conditions, real-time RT-PCR showed very low basal gene expression for TLR2; low for TLR4 and TLR9; moderate for TLR3, TLR5, and TLR7; and undetectable TLR8 (**Figure 2A**). We discovered that, of all the TLRs, only TLR5 gene expression increased significantly after 20-hour stimulation with 33-mer at 250 µM, which contained 33-mer monomers, oligomers, and large thin-plate nanostructures (*p*<0.05) (**Figure 2A**). No such effect was observed at 50 µM (composed of only monomers and oligomers) or after stimulation with 0.068 mg/mL of P-gliadin (**Figure 2B**, **Figure S1**). We used bacterial flagellin from S. typhimurium as a positive control because it is the gold-standard TLR5 agonist (42). As shown in **Figure 2**, Caco-2 cells responded to apically added bacterial flagellin, as previously reported (43), showing an approximately 3.5-fold increase in TLR5 mRNA levels compared with the 33-mer.

Cell viability was assessed using the lactate dehydrogenase (LDH) activity assay, which measures LDH release from cells as a marker of cellular damage. Results showed negligible cellular LDH release across all stimuli, suggesting that the treatment was not harmful to the cells (**Figure 2C**). To assess apoptosis, we determined the ratio of Bak-1 (pro-apoptotic) to Bcl-2 (anti-apoptotic) gene expression (SI, Table S1). No increase in the Bak-1:Bcl-2 ratio was observed for 33-mer stimuli. Regarding P-gliadin, the pro-apoptotic ratio increased 3-fold, suggesting that this gliadin peptide mixture induces mild cellular stress. As reported, P-gliadin stimulation impacts inflammation-related genes (13).

Next, we examined the gene and protein expression of two chemokines, CXCL1 and CXCL8, which are known to be associated with TLR5 activation (42–44). Caco-2 monolayers showed moderate CXCL1 gene expression levels at baseline. After stimulation with the 33-mer structures, a 1.8 ± 0.5-fold increase in gene expression was observed upon stimulation with 250 µM 33-mer, translating to 51.2 ± 3.7 pg/mL protein versus 35.5 ± 2.7 pg/mL for unstimulated wells (*p*=NS) (**Figure 2D**). For P-gliadin, CXCL1 expression increased, although it did not reach significance. At baseline, CXCL8 mRNA expression was low in Caco-2 monolayers, and protein values were below the ELISA detection limit for both 33-mer (data not shown). Data from a previous report showed that 33-mer accumulation did not induce inflammatory markers in Caco-2 cells (13), consistent with our findings. Compared to unstimulated cells, monolayers that were incubated with apical flagellin showed a 4.6 ± 1.5-fold increase in CXCL1 gene expression (*p*<0.05) and a high and significant production of CXCL1 (1205.9 ± 15.3 pg/mL, *p*<0.01) (**Figure 2D**). Likewise, monolayers that were incubated with flagellin showed a 3.4 ± 0.3-fold increase in CXCL8 gene expression and a moderate but significant CXCL8 production (43.5 ± 4.5 pg/mL, *p*<0.01) (**Figure 2E**). Taken together, these results demonstrate that Caco-2 cells respond to apical flagellin stimulation by increasing TLR5 mRNA expression and CXCL1 and CXCL8 production. In contrast, only at the highest 33-mer concentration, at which monomers coexist with oligomers and thin-plate structures, the cells show a moderate but statistically significant increase in TLR5 mRNA expression, without concomitant changes in CXCL1 or CXCL8 levels. These findings suggest that the 33-mer system can modulate TLR5 mRNA expression without eliciting a pro-inflammatory response under the experimental conditions tested.

### The 33-mer peptide colocalizes with TLR5 in Caco-2 cells

Immunostaining of Caco-2 cell monolayers grown on permeable filters was followed by visualization using wide-field deconvolution microscopy, showing TLR5 (green) and 33-mer (red) (**Figure 3A**). We observed that the 33-mer (red) partially overlapped with TLR5 (green), appearing in yellow. The Pearson’s correlation coefficient (PCC) (*p*<0.001) indicates colocalization compared with the control (33, 34) (**Figure 3B**). Transepithelial electrical resistance (TEER) experiments confirmed that the 33-mer at 250 μM does not decrease monolayer permeability (12) (SI; Figure S2), with only 0.06% detected in the basolateral compartment after 20 hours of stimulation (SI, Table S2), consistent with transcellular transport as reported *in vivo* (31,32).

**Figure 3.**
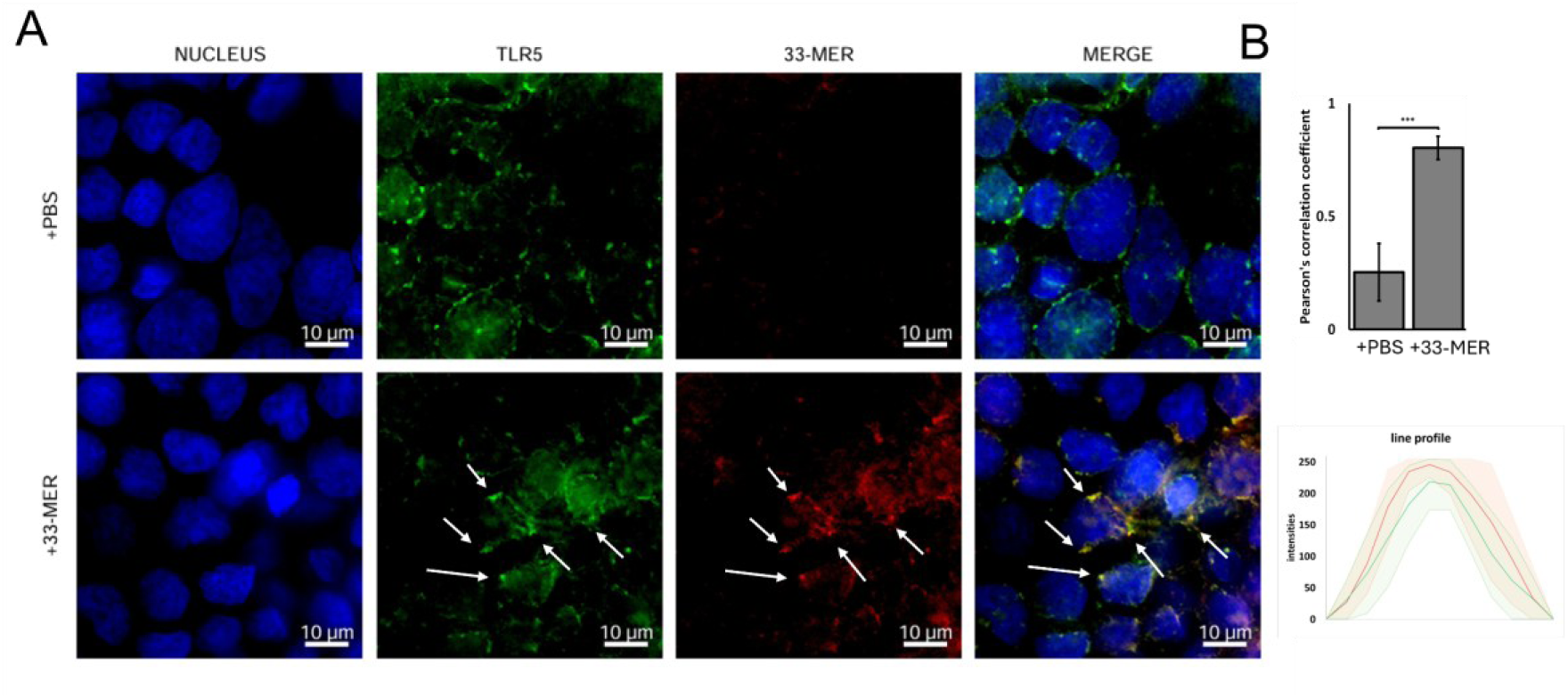
Visualizing the TLR5 and 33-mer in Caco-2 cells. (A) Representative maximum intensity projections of z-stacks spanning 1 µm in thickness from three independent biological experiments (n = 3) are shown. Wide-field fluorescence microscopy images were processed and assembled using OMERO software (33). Immunostaining was performed to visualize TLR5 (green), the 33-mer (red), and nuclei were stained with DAPI (blue). Upon stimulation with 33-mer, partial colocalization with TLR5, shown with arrows, was detected in yellow (MERGE Panel). (B) Pearson’s correlation coefficient (PCC) was calculated from 12 regions of interest per condition (34), revealing differences in colocalization between 33-mer-treated and PBS-treated cells (p < 0.001).

Taken together, these findings demonstrate transcellular transport of the 33-mer in Caco-2 cells and reveal its partial colocalization with TLR5, suggesting spatial proximity between the peptide and the receptor.

### The 33-mer peptide does not directly activate TLR5 but attenuates flagellin-induced TLR5/NF-κB signaling

Next, we investigated TLR5-dependent NF-κB activation triggered by 33-mer nanostructures using the HEK-Blue-hTLR5 cell line, which overexpresses TLR5 and its parental HEK-Blue-Null cell line as a control. The signal produced after stimulation with flagellin was normalized to 100 % (full activation), and the signals from the other stimuli were expressed as a percentage of the flagellin response. Addition of neutralizing anti-hTLR5 mAb 1 hour before flagellin stimulation reduced the signal to 25 ± 2.2%, close to the basal level (15,7 ± 0.1%). No NF-κB activation was detected after stimulation with increasing concentrations of 33-mer or P-gliadin mediated by TLR5 (**Figure 4A**). In other contexts, such as neurodegenerative diseases, the formation of peptide oligomers and superstructures is a hallmark of protein aggregation-related or amyloid diseases, such as Aβ aggregation in Alzheimer’s disease (45). It is known that Aβ peptide and its oligomers are able to modulate the expression and activation of TLRs (46–49), in particular TLR4 and TLR5, similar to amyloid-like proteins present in bacterial biofilms (35,44). In particular, oligomeric and fibrillar Aβ attenuate flagellin-mediated activation of human TLR5 without activating TLR5 signaling on its own (46). Considering some structural similarities between 33-mer and Aβ oligomers (50, 51), we investigated whether the interaction between the 33-mer and TLR5 could neutralize flagellin-mediated receptor activation, akin to a neutralizing anti-TLR5 antibody. Consequently, we pre-incubated the cells for 1 hour with increasing concentrations of 33-mer (25-250 µM), followed by the addition of flagellin. **Figure 4B** shows that the 33-mer peptide was capable of dose-dependent inhibition of flagellin-induced NF-κB activation, starting at 50 µM with a reduction of the signal to 71.4 ± 3.6 % (*p*=NS) and reaching significance at 150 µM and 250 µM with reduced values of 64.7 ± 5.3 % (*p*<0.05) and 43.5 ± 3.2 % (*p*<0.01), respectively. These data suggest that the greater decrease in flagellin-induced NF-κB activation at higher 33-mer concentrations is due to the 33-mer supramolecular system interfering with the TLR5 activation pathway. To test whether this inhibition is specific to the 33-mer, we used the metabolic 33-mer DGP as a second control; it has triple point mutations at Q10E, Q17E, and Q24E and forms only small oligomers at these concentrations (12). Dose–response experiments revealed that 33-mer DGP neither induced TLR5-mediated NF-κB activation on its own nor inhibited flagellin-induced activation at increasing concentrations (**Figure 4C-D**).

**Figure 4.**
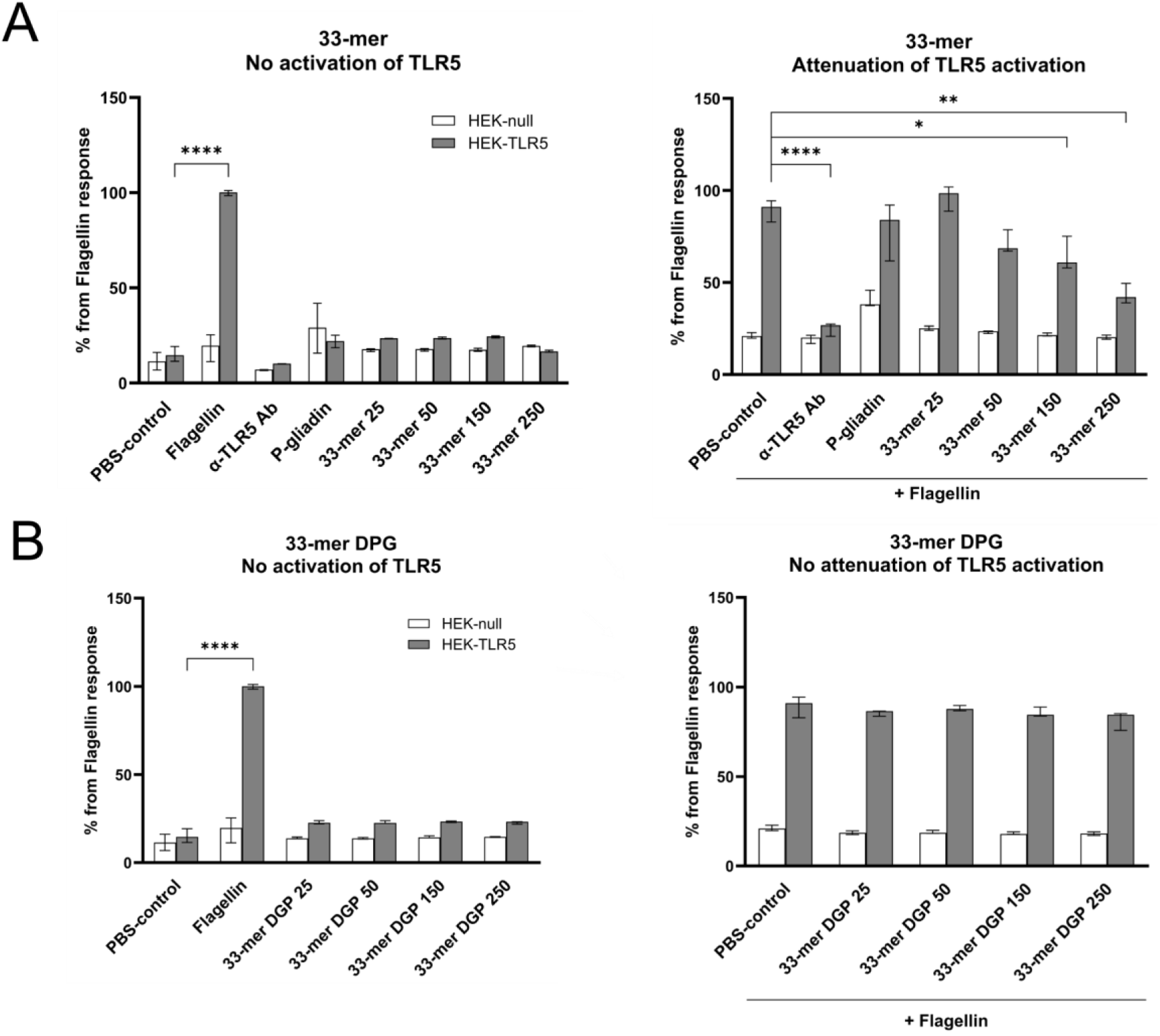
33-mer does not activate the NF-κB pathway mediated by TLR5, but is capable of mitigating TLR5-mediated NF-κB activation by flagellin. (A) Left: Ultrapure Flagellin (10 ng/mL) was used as the positive control for TLR5-mediated NF-κB activation. Values were normalized to 100% (*p*<0.0001). The neutralizing antibody itself showed no effects. Increasing concentrations of 33-mer or 33-mer DGP (25 to 250 µM) and 0.068 mg/mL P-gliadin did not activate NF-κB (P=NS). Right: Cells were pre-incubated for 1 hour with PBS, neutralizing anti-TLR5 Ab, P-gliadin, or increasing concentrations of 33-mer or 33-mer DGP and subsequently, flagellin was added for 20 hours. Data revealed that PBS control/flagellin induced NF-κB activation (89 ± 3 %), and that blocking with the neutralizing anti-TLR5 Ab strongly inhibited flagellin-induced NF-κB activation to 24.9 ± 2.1 % (*p*<0.0001). P-gliadin did not show inhibitory effects on flagellin-induced NF-κB activation (79.3 ± 9.1 %, P=NS), whereas 33-mer significantly reduced flagellin’s activation of TLR5-mediated NF-κB activation at 33-mer 150 µM (64,7 ± 5.3 %, *p*<0.05) and 250 µM (43.5 ± 3.2 %, *p*<0.01). (B) No effect was observed for the control peptide 33-mer DGP under the same experimental conditions. Values are given as mean ± interquartiles (n=6).

Taking into account these results, we demonstrated that the 33-mer system, composed of monomers, oligomers, and thin-plates structures, interacts with TLR5 in a way that the 33-mer DGP peptide and its oligomers can not. In fact, the behavior of 33-mer is similar to that observed for oligomeric and fibrillar Aβ (46). Finally, in line with the previously reported structural and functional similarities between gliadin peptides and pathogen-derived molecules (52–54), our finding that the 33-mer attenuates flagellin-induced TLR5 activation adds a new layer of complexity by suggesting a potential receptor-mediated link that warrants further investigation

### A hypothetical mechanistic model of the TLR5/33-mer interaction by molecular docking

To gain structural insight into our experimental observations, we employed molecular docking as an initial exploratory approach to assess the structural complementarity and potential molecular interactions of the monomeric 33-mer with TLR5, using the 33-mer DGP as a comparator, as the latter did not attenuate flagellin-induced TLR5 signaling in our functional assays. For this purpose, we performed blind molecular docking trials using the TLR5 structure obtained from the crystal structure of the TLR5 homodimer (TLR5A/TLR5B) in complex with a fragment of Salmonella flagellin (**Figure 5A, left**) (36). The reported crystal structure of the complex consists of a 2:2 symmetric heterodimer complex of the N-terminal, which comprises nearly two-thirds of the entire TLR5 horseshoe-shaped ectodomain, with a fragment deletion of flagellin (ΔD0) (36). The ectodomain contains leucine-rich repeats (LRRs) that serve as the ligand-binding site. Only the conserved D1 domain of the flagellin ΔD0 fragment forms non-covalent interactions with the concave surface of TLR5 chain A (36). Importantly, although flagellin monomers polymerize to form the bacterial flagellar filament, the available TLR5– flagellin crystal structures describe the interaction with monomeric flagellin (36) that promotes receptor dimerization, thereby activating NF-κB-dependent inflammatory signaling (20, 55–57)(**Figure 5B**). By analogy, although the 33-mer peptide can assemble into higher-order structures, we hypothesize that TLR5 recognition is primarily mediated by the monomeric form of the peptide. Docking results show that, in the best-scored output of the TRL5/33-mer docking system (**Figure 5A, right**), the monomer of the 33-mer peptide (red) accommodates the entire concave surface of TLR5 and a significant part of the TLR5 dimerization region. The 33-mer forms multiple polar and non-polar interactions with TLR5 (**SI, Table S3a**). The extent of the area involved in the TLR5 dimerization region was calculated as 652.6 Å^2^ (crystal structure), of which 348.7 Å^2^ (53.4 %) is covered by the 33-mer peptide (docking output).

**Figure 5.**
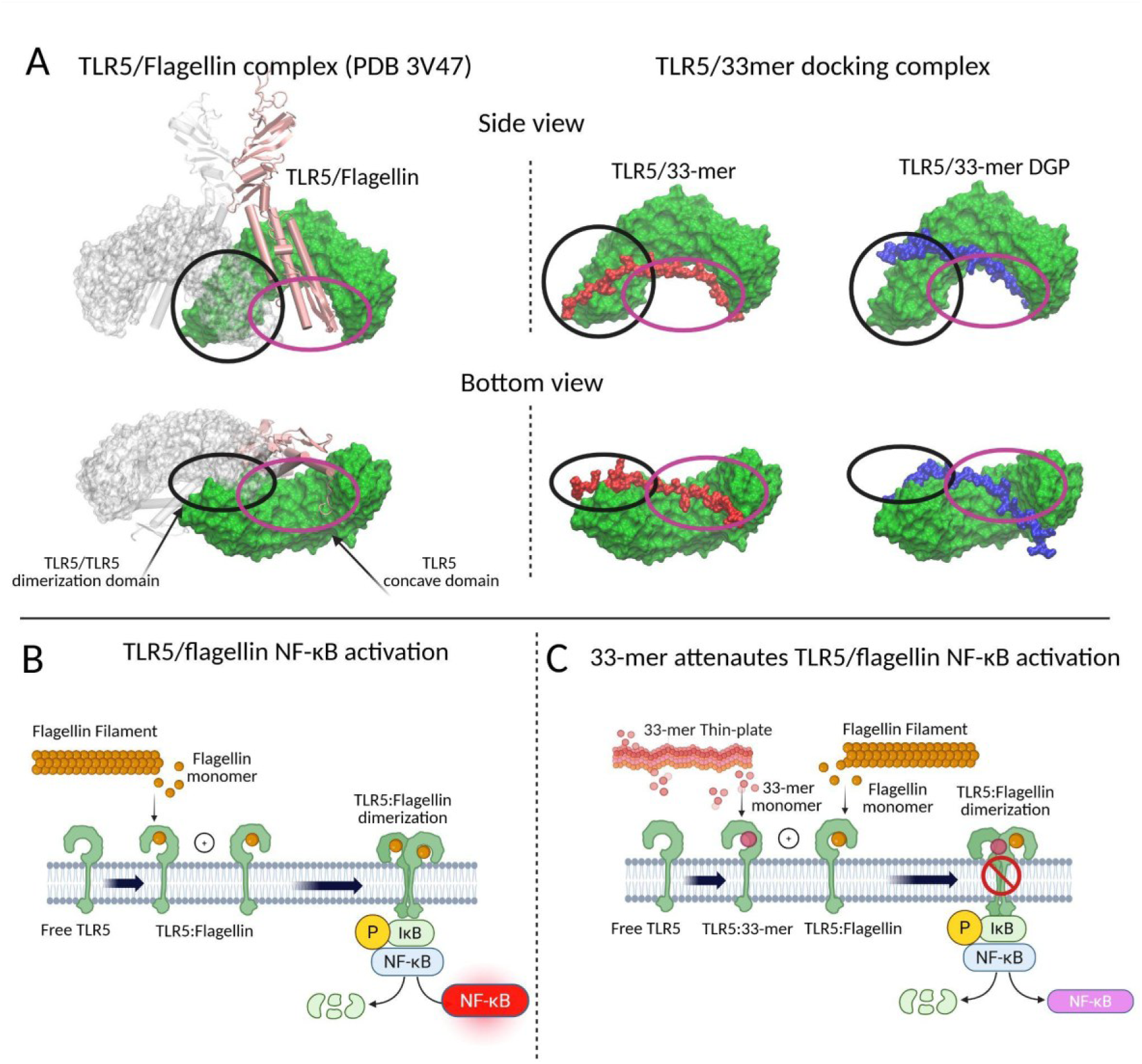
Hypothetical interaction of the 33-mer peptides with TLR5. (A) Left: Chain A of the crystallographic structure of TLR5 in complex with Salmonella flagellin was used as a model (PDB ID 3V47); chain B is shown in cartoon representation (grey) as a reference but was not used in the protein-peptide docking. Right: The top-ranked docking conformations of monomeric 33-mer (red) and 33-mer DGP (blue) with chain A of TLR5 are shown in side and bottom views. The concave surface (black circle) and dimerization region (purple circle) of TLR5 are highlighted. The corresponding docking scores were −280.20 for the TLR5–33-mer complex and −293.07 for the TLR5–33-mer DGP complex. Docking scores are dimensionless and should not be interpreted as binding free energies. More information in SI. (B) Schematic of TLR5 activation mediated by the interaction of monomeric flagellin, which induces TLR5 dimerization and downstream pathway activation. (C) Hypothetical attenuation mechanism of flagellin-mediated TLR5 activation, due to 33-mer/TLR5 interaction that impacts the TLR5-TLR5 dimerization domain. Created in BioRender. Dodero, V. (2026).

Taking into consideration that 33-mer DGP did not exert an effect on the activation or mitigation of flagellin/TLR5 NF-κB pathway activation (**Figure 4 C-D**), we ran a second docking experiment using the model of 33-mer DGP (19). In comparison, the best docking pose of the TLR5/33-mer DGP complex shows that the 33-mer DGP is not fully located in the dimerization domain of TLR5 chains, in part guided by electrostatic interactions with acidic residues of the 33-mer DGP peptide (**SI, Figure S2 and Table S3b**). The 33-mer DGP peptide occupied an area of 222.5 Å², corresponding to 34.1% of the TLR5 dimerization interface, which was lower than the 54% occupied by the 33-mer peptide. Interestingly, the docking models suggested different binding modes, with the 33-mer showing a greater predicted overlap with the TLR5 dimerization interface than 33-mer DGP. Based on these observations, we propose a hypothetical structural mechanism (**Figure 5C**) in which 33-mer thin-plate nanostructures may act as reservoirs of monomeric peptides, akin to flagellin assembled in the filament, potentially increasing their local concentration for interaction with TLR5. This plausible binding attenuates flagellin-induced receptor activation, providing an initial structural rationale for our functional observations and establishing a testable hypothesis for future mechanistic investigations.

### Limitations of the study

Several limitations of the present study should be acknowledged. First, although the 33-mer increased TLR5 mRNA expression in Caco-2 monolayers, TLR5 protein abundance and cell-surface expression were not quantitatively assessed. Second, attenuation of flagellin-induced TLR5/NF-κB signaling was demonstrated using a reporter-cell system stimulated with purified S. Typhimurium flagellin, which does not reproduce the complexity of physiological intestinal responses to bacteria. Third, although partial colocalization and molecular docking suggest a possible association between the 33-mer and TLR5, no direct binding, competition with flagellin, or interference with receptor dimerization was performed and therefore remains experimentally unverified. Finally, because the 33-mer system is composed of monomers, oligomers, and thin-plates assemblies, the observed effects cannot be attributed to a specific 33-mer nanostructure, but only to the 33-mer system.

### Potential clinical implications and future directions

TLR5 is expressed by intestinal epithelial and immune cells and contributes to intestinal immune surveillance and mucosal homeostasis by recognizing bacterial flagellin (20, 35, 42, 58). In this context, our findings raise the possibility that the accumulation and supramolecular organization of the 33-mer gliadin peptide may influence intestinal responses to bacterial stimuli. The selective upregulation of TLR5 mRNA in Caco-2 monolayers without inflammation, together with attenuated flagellin-induced TLR5 activation in reporter cells, provides a rationale for further investigating the potential interplay between gliadin peptide assemblies and microbial recognition pathways. Given the established role of the proteolysis-resistant peptides in gluten-related disorders and in particular the 33-mer gliadin peptide in CeD, our findings revealing properties beyond its immunogenicity may have broader implications for understanding the early triggers of GRDs, including CeD and non-celiac gluten/wheat sensitivity (NCGS/NCWS) (4,6), and their association with gastrointestinal disorders, including irritable bowel syndrome (IBS) and Crohn’s disease (59,60). Interestingly, antibodies against bacterial flagellin have been detected in patients with Crohn’s disease (61) and NCWS (62). Therefore, future studies should investigate the effects of 33-mer nanostructures in more physiologically relevant models, including human intestinal organoids and patient-derived intestinal samples, where it will be possible to consider the influence of epithelial polarity and its compartmentalization. These approaches may help to disclose the contribution of 33-mer supramolecular organization to TLR5-mediated responses and determine its potential relevance to intestinal immune homeostasis and gluten-related disorders.

## Conclusions

We found that apical stimulation of Caco-2 monolayers with the 33-mer system increased TLR5 mRNA expression without inducing CXCL8 production or significantly increasing CXCL1 levels. In contrast, flagellin from S. Typhimurium increased TLR5 mRNA expression and significantly increased chemokine production, consistent with an inflammatory response. Interestingly, immunofluorescence microscopy revealed partial colocalization between the 33-mer and TLR5, suggesting spatial proximity. Using reporter cells, we established that 33-mer does not activate TLR5 on its own. But pre-incubation with 33-mer followed by flagellin stimulation resulted in a concentration-dependent attenuation of TLR5/NF-κB signaling, an effect not observed with the control peptide 33-mer DGP. Molecular docking suggested a possible structural mechanism in which the 33-mer could occupy the concave region of TLR5 and part of its dimerization interface, potentially interfering with receptor activation. Together, we hypothesize that 33-mer supramolecular assemblies may act as reservoirs of monomeric peptides, potentially increasing their local availability to modulate flagellin-induced TLR5 signaling. Further studies are in progress to test this hypothesis and elucidate the contributions of distinct 33-mer supramolecular species to TLR5-mediated innate immune signaling.

## Acknowledgments

This work was funded by the Deutsche Forschungsgemeinschaft (DFG), grant number 430578458. Initial results were obtained during the V.I.D. Alexander von Humboldt fellowship (HERMES). The authors thank Dr. Philipp Dörfler for experiments on reporter cells and microscopy and Mrs. Carmela Michalek for cell culture support. We acknowledge support for the publication costs by the Open Access Publication Fund of Bielefeld University and the Deutsche Forschungsgemeinschaft (DFG).

## Contributions

V.I.D. conceived, supervised the project and secured funds; K.M.L. and V.I.D. designed and performed the stimulation experiments; K.M.L. performed and analyzed qPCR and ELISA; W.H. and V.I.D. performed and analyzed wide-field fluorescence microscopy experiments; V.S. and C.S. conducted 33-mer ELISA experiments; D.S.V. performed, interpreted, and analyzed computational experiments; J.H., S.M., C.S., T.H., and V.I.D. provided resources; K.M.L., D.S.V., and V.I.D. wrote the manuscript. All the authors reviewed and approved the final version of the manuscript.

## Ethics declarations

The authors declare no competing financial interests.

