## Supplementary material for "The immunogenic 33-mer gliadin peptide attenuates flagellin-induced TLR5/NF-κB signaling": SI

### 1. Toll-Like Receptor mRNA expression promoted by P-gliadin stimulation

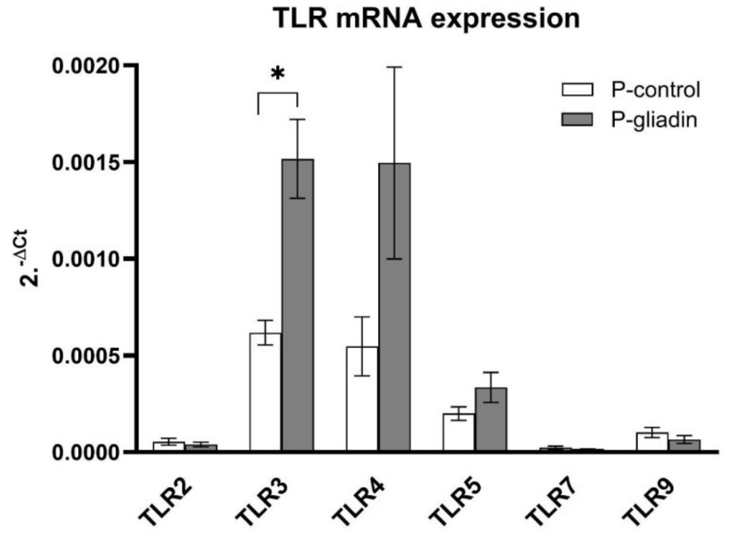

**Figure S1. TLR gene expression by P-gliadin (0.064 mg/mL) in Caco-2 monolayers.** Screening of TLR expression in Caco-2 cells by P-gliadin revealed low baseline expression of TLR2, TLR7 and TLR9 and moderate expression of TLR3, TLR4, and TLR5. Only TLR3 expression was increased significantly after stimulation ($p$<0.05).

### 2. Cell viability

Trypan blue exclusion was performed to assess cell viability, revealing that the cells were viable after treatment (Table S1a). The lactate dehydrogenase (LDH) release measured in $H_2O$-lysed cells was taken as the 100% value and as the control. The results were expressed as milliunits of LDH activity per milligram of protein assayed by the bicinchoninic acid assay. No significant differences in LDH release were observed between unstimulated and treated samples, indicating that no cell lysis had occurred. To assess apoptosis, the ratio of Bak-1 (pro-apoptotic) to Bcl-2 (anti-apoptotic) gene expression was determined. As a moderate positive control for the induction of apoptosis in Caco-2 cells, sodium butyrate (20 mM) (1) was added to parallel monolayers (Table S1b).

**Table S1a**. Trypan Blue results.

| Stimuli | Trypan Blue exclusion (%) |
|---|---|
| Unstimulated | 87 ± 3 |
| 33-mer 50 µM | 90 ± 3 |
| 33-mer 250 µM | 89 ± 6 |

**Table S1b**. Apoptosis results.

| Stimuli | Ratio Bak1/Bcl2 |
| --- | --- |
| Unstimulated | 1 |
| 33-mer 50 μM | 1.18 |
| 33-mer 250 μM | 1.13 |
| P-Gliadin | 1.13 |
| Flagellin | 0.86 |
| Sodium Butyrate 20 mM | 1.45 |

### 3. TEER measurement

TEER values were measured in Ohms using a Millicell Electrical Voltohmmeter (EVOM) with chopstick electrodes. After the cells had grown adherent and confluent, the TEER value was measured regularly without touching the cell layer. Cells were ready for stimulation after 21 days of culture. The electrodes were inserted vertically into the apical and basolateral compartments simultaneously, and the stable value was determined with EVOM. Following the same procedure, the values of one insert without cells (RBLANK) and inserts with monolayers were determined (RTOTAL). Subsequently, the blank value RBLANK was subtracted from the measured RTOTAL, and the result was multiplied by the area (MAREA) on which the cells were grown. Only wells with initial values above 450 $\Omega cm2$ were employed.

$$R_{TISSUE}(\Omega) = R_{TOTAL} - R_{BLANK} \quad \text{(Eq. 1)}$$

$$TEER\ REPORTED = R_{TISSUE}(\Omega) \times M_{AREA}(cm^2) \quad \text{(Eq. 2)}$$

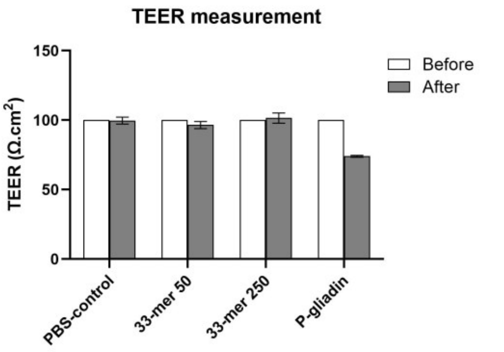

**Figure S2.** TEER measurements before and after 20 h of stimulation with phosphate-buffered saline (PBS), 33-mer (50 μM and 250 μM), and P-Gliadin (0.064 mg/mL)

### 4. Quantification of the 33-mer gliadin peptide basolateral by ELISA

The presence of 33-mer superstructures in apical and basolateral solutions after 1 hour and 20 hours of stimulation was extracted by mixing 10 µL of the sample with 90 µL of 60 % (vol:vol) ethanol in a rotary shaker for 1 hour at RT. Then, the suspension was centrifuged at 2500 × g for 10 minutes, and the supernatant was collected and used for quantification.

The relative amounts of the 33-mer immunogenic peptides in apical and basolateral solutions were quantified by a G12 mAb competitive ELISA (2-5). Maxisorp microtitre plates (Nunc, Roskilde, Denmark) were coated with PWG gliadin solution (Sigma-Aldrich, USA) and incubated for 2.5 hours at 37 ºC. Afterward, the plates were washed with PBS containing 0.05 % Tween 20 and then blocked with 3 % bovine serum albumin (BSA) (A2153; Sigma-Aldrich) in PBS for 1 hour at RT. Serial dilutions of the 33-mer peptide from apical and basolateral samples were made, to each of which a solution of HRP-conjugate with G12 mAb was added. The samples were pre-incubated for 1 hour at RT and spun down at 300 RPM. Then, the plates were washed, and substrate peroxidase solution (Tetramethylbenzidin (TMB), Sigma, USA) was added. The reaction was stopped with 1 M sulfuric acid, and the absorbance at 450 nm was measured (UVM340 microplate reader; Asys Hitech GmbH, Eugendorf, Austria). Two separate assays were performed, each with two repetitions. The calculations were performed according to the manufacturer's instructions. A quantification limit of <0,0001 mg/ml was determined by the G12 mAb ELISA assay (Figure S3).

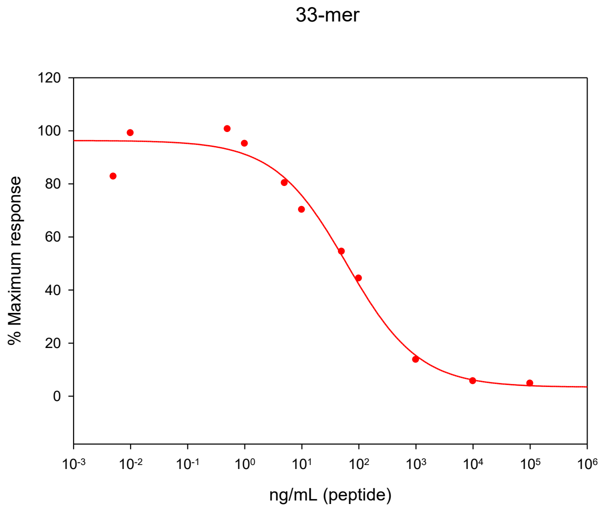

**Figure S3**. A quantification limit of < 0.0001 mg/mL was determined using the G12 mAb ELISA assay.

**Table S2**. Recovery of the 33-mer peptide after stimulation obtained by ELISA.

| Stimulation time | Solutions | Concentration 33-mer (mg/ml) | Recovery % |
|---|---|---|---|
| | stimulation | 0,96 | 100 |
| 20 hours | apical 1 | 0.76 ± 0.07 | 76.00 |
| | basolateral 1 | 0,0006 | 0,06 |
| | apical 2 | 0.80 ± 0.04 | 80.00 |
| | basolateral 2 | 0,0004 | 0,04 |
| 1 hour | apical 3 | 0.74 ± 0.01 | 74.00 |
| | basolateral 3 | <0,0001 | <0,0001 |
| | apical 4 | 0.69 ± 0.1 | 69.00 |
| | basolateral 4 | <0,0001 | <0,0001 |

### 5. Co-localization analysis

Using the ImageJ plugin Coloc 2, it is possible to analyze the co-localization of fluorescence signals from two channels recorded with fluorescence microscopy. Therefore, the single-channel images must undergo deconvolution to assign the signals to the correct z-levels, and background noise must be removed by adjusting the intensities in a linear fashion. Specifying a region of interest (ROI) in one of the images allows selection of the area where the co-localization analysis shall be performed. The output of the Coloc 2 plugin includes a 2D histogram. Pearson's correlation coefficient (PCC) measures the overlap between the pixel values of two images and ranges from [-1,1] (-1: negative correlation, 0: no correlation, 1: positive correlation) (6). Twelve representative areas were outlined using the freehand tool and Coloc 2 was used to analyze the co-localization of the two signals across the stack of images. The PCC tending towards 1 indicates a highly linear relationship between the pixels of the two images.

### 6. Molecular docking

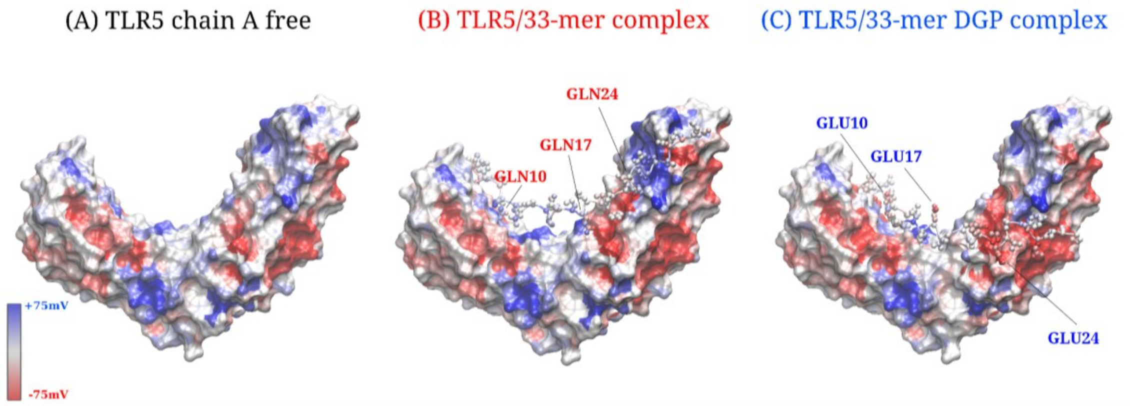

**Figure S4. Charge distribution function of the TLR5/33-mer complexes.** The charge distribution function is shown for TLR5 chain A (A), for the TLR5/33-mer complex (B), and TLR5/33-mer DGP complex (C). The residues implicated in the deamidation of the 33-mer DGP are shown. The analysis was performed using the Protein-Sol server (https://protein-sol.manchester.ac.uk/) (7).

**Table S3a**. Contacts between TLR5 and the 33-mer peptide of the best-scoring model. The analysis was performed using an *ad hoc* Biopython script (8). A cutoff of 3.5 Å was set to calculate contacts. The SASA was calculated using the Shrake-Rupley algorithm.

| Aminoacid TLR5 | Residue number | SASA % | Amino acid 33-mer | Residue number | SASA % | Distance (Å) |
|---|---|---|---|---|---|---|
| TYR | 30 | 57.91 | GLN | 2 | 73.24 | 2.36 |
| TYR | 51 | 19.42 | PRO | 5 | 18.1 | 2.99 |
| TYR | 51 | 19.42 | PHE | 6 | 104.93 | 3.34 |
| GLN | 74 | 63.82 | PRO | 5 | 18.1 | 2.56 |
| ILE | 100 | 25.36 | GLN | 8 | 49.41 | 3.18 |
| LYS | 102 | 13.35 | PRO | 9 | 47.49 | 2.85 |
| VAL | 124 | 12.08 | GLN | 8 | 49.41 | 2.68 |
| LYS | 301 | 42.7 | GLN | 17 | 84.58 | 3.44 |
| GLN | 325 | 22.91 | GLN | 17 | 84.58 | 1.7 |
| GLN | 349 | 36.61 | PRO | 19 | 52.69 | 2.68 |
| GLN | 349 | 36.61 | LEU | 18 | 156.93 | 3.48 |
| GLN | 349 | 36.61 | GLN | 17 | 84.58 | 3.34 |
| TYR | 373 | 95.84 | PRO | 19 | 52.69 | 3.06 |

| | | | | | | |
|---|---|---|---|---|---|---|
| HIS | 375 | 42.97 | GLN | 22 | 62.35 | 3.48 |
| GLN | 399 | 27.67 | PRO | 23 | 64.97 | 3.01 |
| GLN | 399 | 27.67 | GLN | 22 | 62.35 | 3.3 |
| LYS | 401 | 105.74 | PRO | 28 | 45.75 | 3.29 |
| SER | 452 | 11.15 | TYR | 27 | 84.19 | 2.74 |
| SER | 452 | 11.15 | PRO | 28 | 45.75 | 2.54 |
| GLY | 453 | 54.34 | PRO | 30 | 78.46 | 3.02 |

**Table S3b**. Contacts between TLR5 and the 33-mer-DGP peptide of the best scored model. The analysis was performed using an *ad hoc* Biopython script (8). A cutoff of 3.5 Å was set to calculate contacts. The SASA was calculated using the Shrake-Rupley algorithm.

| Aminoacid TLR5 | Residue number | SASA % | Amino acid 33-mer-DGP | Residue number | SASA % | Distance (Å) |
|---|---|---|---|---|---|---|
| GLN | 74 | 20.78 | PRO | 5 | 122.21 | 3.38 |
| PHE | 75 | 20.53 | PRO | 7 | 46.87 | 2.66 |
| ILE | 99 | 22.94 | PHE | 6 | 44.39 | 2.74 |
| GLU | 123 | 34.29 | PHE | 6 | 44.39 | 3.47 |
| PHE | 180 | 52.18 | PRO | 16 | 61.16 | 2.09 |
| ARG | 203 | 38.6 | PRO | 12 | 12.6 | 2.58 |
| ASN | 265 | 23.9 | LEU | 18 | 66.84 | 3.48 |
| PHE | 273 | 21.46 | GLN | 29 | 94.45 | 3.27 |
| HIS | 275 | 84.53 | GLU | 24 | 125.91 | 3.12 |
| LYS | 303 | 50.92 | PRO | 21 | 10.87 | 1.96 |
| PHE | 305 | 33.81 | GLN | 29 | 94.45 | 2.73 |
| GLN | 325 | 46.47 | TYR | 20 | 87.75 | 2.82 |
| ASN | 329 | 20.36 | GLN | 29 | 94.45 | 2.72 |
| ASN | 329 | 20.36 | PRO | 28 | 20.78 | 3.42 |
| GLN | 349 | 80.92 | TYR | 20 | 87.75 | 3.46 |
| PHE | 351 | 18.11 | PRO | 23 | 28.01 | 2.93 |
| SER | 354 | 10.19 | GLN | 31 | 92.73 | 2.47 |
| ARG | 377 | 112.84 | TYR | 27 | 116.15 | 2.94 |
